# Specific origin selection and excess functional MCM2-7 loading in ORC-deficient cells

**DOI:** 10.1101/2024.10.30.621095

**Authors:** Yoshiyuki Shibata, Mihaela Peycheva, Etsuko Shibata, Daniel Malzl, Rushad Pavri, Anindya Dutta

## Abstract

The six subunit Origin Recognition Complex (ORC) loads excess MCM2-7 on chromosomes to promote initiation of DNA replication and is believed to be important for origin specification. Mapping of origins in cancer cell lines engineered to delete three of the subunits, *ORC1*, *ORC2* or *ORC5* shows that specific origins are still used and are mostly at the same sites in the genome as in wild type cells. The few thousand origins that were up-regulated in the absence of ORC suggest that GC/TA skewness and simple repeat sequences facilitate, but are not essential for, origin selection in the absence of the six-subunit ORC. Despite the lack of ORC, excess MCM2-7 is still loaded at comparable rates in G1 phase to license dormant origins and is also repeatedly loaded in the same S phase to permit re-replication. Thus, origin specification and excess MCM2-7 loading on origins do not require the six-subunit ORC in human cancer cell lines.

## INTRODUCTION

The six-subunit ORC is essential for DNA replication in eukaryotes because it helps load the double hexameric MCM2-7 on chromatin and this MCM2-7 hexamer forms the core of the replicative helicase that unwinds origins of replication and initiates DNA replication (1–5). The loading of the MCM2-7 double-hexamer around DNA is also called licensing of origins or formation of the pre-Replicative complex (preRC). Because ORC is among the first replication initiation factors to bind DNA and then help load MCM2-7 on DNA, ORC plays an important role in specifying the locations of origins in most eukaryotes. ORC also loads an excess of MCM2-7 on chromatin relative to what is required to fire the normal complement of ∼50,000 origins per cell cycle, and this MCM2-7 excess is critical for the firing of dormant origins when replication is interrupted by the premature termination of replication forks emerging from the first set of origins (6,7). Thus, this dormant origin firing is a measure of whether excess functional MCM2-7 is loaded in the cells. Finally, the licensing and re-licensing of MCM2-7 on origins when certain cell-cycle pathways are de-regulated, leads to re-replication of the genome, and this is another measure of the robustness of functional MCM2-7 loading (8–11).

There is solid evidence that ORC is essential for licensing origins in yeasts, flies and mammalian cells and *in vitro* (2,3,12–19). In addition, hypomorphic mutations in ORC subunits lead to microcephalic primordial dwarfism resembling Meier-Gorlin syndrome (20–22). In this syndrome, all tissues are not affected, but specific tissues like the patella, ears and the skeleton are smaller than normal. This phenotype is interpreted as consistent with an essential role of ORC in DNA replication and cell proliferation, though it is difficult to explain why a mutation in a complex essential for DNA replication and cell proliferation globally has such tissue-specific effects.

On the other hand, we have generated cancer cell lines in which one of the ORC subunits, ORC1, ORC2 or ORC5 is mutationally inactivated by CRISPR-Cas9 and the cells are still viable and proliferate with DNA replication (23,24). The mutational inactivation results in undetectable levels of ORC1 and ORC5 in the *ORC1Δ* and *ORC5Δ* cells, respectively, and although 0.23% of a truncated ORC2 (∼350 molecules/cell) was reported in the *ORC2Δ* cells (25), we will address this later. We are thus faced with a paradox: while ORC is required for DNA replication in most instances, how do these cancer cell lines replicate and survive without the six-subunit ORC? One possibility is that in the absence of ORC an imprecise loading of MCM2-7 leads to the licensing of diffuse, non-specific origins of replication, so that it would be impossible to map precise origins of replication in a population of cells. Also, the MCM2-7 loading could be so inefficient that dormant origin licensing or repeated origin licensing in the same S phase are not possible. It is worth noting that some of the cell lines retain different subcomplexes of ORC on chromatin. In the *ORC2Δ* cells ORC1 and ORC6 alone are on the chromatin, in the *ORC1Δ* cells, ORC2-6 load on chromatin at near wild type levels, and the *ORC5Δ* cells allow ORC1, 2, 3 and 6 to associate with chromatin at near wild type levels (23,24). Thus, another possibility is that different subcomplexes of ORC specify different subsets or origins and have differing ability to load excess MCM2-7.

We therefore decided to stringently test whether, (1) no origins or a different set of origins are selected in the ORC knockout cells or (2) a highly compromised licensing machinery without ORC can load enough MCM2-7 at a reasonable rate in G1 for replication from dormant origins or for re-replication in S phase.

## MATERIALS AND METHODS

### Cell lines and cell culture

HCT116 p53−/− cells (26) is a generous gift from Fred Bunz, Johns Hopkins. Cells were maintained in McCoy’s 5A-modified medium supplemented with 10 % fetal bovine serum. HCT116 p53-/- *ORC2Δ* (also referred to as ORC2KO), *ORC1Δ* (ORC1KO), or *ORC5Δ* (ORC5KO) cells were generated using CRISPR-Cas9 technologies (23,24).

### Subcellular fractionation and Western blots

Chromatin fractionation was performed as previously described (27). In brief, Cells were lysed in buffer A (10mM HEPES [pH7.9], 10 % glycerol, 1mM DTT, Protease inhibitor cocktail) with 0.1% Triton X100, incubated for 8 min on ice and centrifuged at 1300g for 5min. Pellets were dissolved in buffer B (3 mM EDTA, 0.2 mM, EGTA, 1 mM DTT, protease inhibitor cocktail) and incubated for 30 min on ice. Cell lysates were centrifuged at 1700g for 5 min and supernatants were removed. Pellets were lysed in 2 x SDS sample buffer and sonicated (P3).

Antibodies used were as follows. ORC1(4731, Cell Signaling Technology), ORC2 (sc-32734, Santa Cruz Biotechnology or generous gift from Bruce Stillman, Cold Spring Harbor Laboratory), ORC3 (sc-23888, Santa Cruz Biotechnology), ORC5 (28), HSP90 (sc-13119, Santa Cruz Biotechnology), LaminA/C (2032, Cell Signaling Technology), MCM2 BM28 (610700, BD), MCM7 (sc-9966, Santa Cruz Biotechnology), MCM6 (sc-9843, Santa Cruz Biotechnology).

### Immunoprecipitation

Cells were lysed in IP buffer (50 mM Hepes, 150 mM NaCl, 1mM EDTA, 2.5 mM EGTA, 10 % Glycerol, 0.1 % Tween20) and precleared by centrifuge, then cell lysate was incubated with ORC3 serum (29) bound Dynabeads Protein G (10004D, Fisher Scientific) for 18 h at 4°C and then eluted with 2 x SDS sample buffer.

### Chromatin flow cytometry

The method has been described in detail in (30). Anti- MCM2 BM28 (610700, BD), anti-Mouse IgG secondary antibody, Alexa Fluor™ 488 (A-11029, Thermo Fisher), DAPI (32670, Millipore Sigma) were used. Fluorescence was measured using a BD LSRFortessa and data was analyzed using Flow Jo. DAPI with secondary antibody alone was used to determine MCM2 positive population.

### Ergodic rate analysis

The method has been described in detail in (30) except Flow Jo. was used to export G1- MCM^DNA^ (number of cells with MCM loaded on DNA in G1 phase) raw data to CSV file (Supplementary **Table S4**). G1-MCM^DNA^ raw data were divided into 10 equal sized bins between the lowest and highest MCM value and used to calculate MCM loading rate. For a mean MCM loading rate, all 10 MCM loading rates were averaged.

The MCM loading rate was calculated using MATLAB (MathWorks). The rate calculation was

W_n_ (MCM loading rate in bin N) = α(2-F)/f_n_

α = L_n_(2)/doubling time

F = number of G1-MCM^DNA^ cells/total number of cells in sample

f_n_ = number of cells in bin n/number of G1-MCM^DNA^ cells

Sample MATLAB code was as follows

alpha_WT= log(2)/19.5;

F_WT_1 = 0.288143;

%calculate lowest and highest MCM values%

maxMCM_WT_1 = max(WT_1(:,1));

minMCM_WT_1 = min(WT_1(:,1));

%create histogram with 10 bins and specified first and last bin limits%

h10_WT_1 = histogram(WT_1(:,1), ‘NumBins’, 10, ‘BinLimits’, [minMCM_WT_1, maxMCM_WT_1]);

%calculate fn within each bin%

totalf_WT_1 = size(WT_1);

fn10_WT_1=(h10_WT_1.Values)/totalf_WT_1(1,1);

%calculate mean w%

w10mean_WT_1 = mean(alpha_WT.*(2- F_WT_1)./fn10_WT_1)

### Hydroxyurea sensitivity and molecular combing

2500 (WT, ORC1KO, ORC2KO) or 10,000 (WT, ORC5KO) cells were seeded into 6-well plates the day before addition of the indicated amounts of hydroxyurea. After 5 days (WT, ORC1KO ORC2KO) or 7 days (WT, ORC5KO), cells were fixed and stained in a solution of 20% formaldehyde, 80% methanol, and 0.25% crystal violet, and rinsed with deionized water. Images were quantified with Image J.

For molecular combing assay, cells were treated with 200 μM of hydroxyurea for 4 h, then pulse- labeled with 100 μM 5-chlorodeoxyuridine (ldU) followed by 250 μM 5- iododeoxyuridine (CIdU) for 30 min each. 1M Cells suspended in trypsin were mixed with 1.2% melted low agarose and transferred into plug mold (170-3713, Bio-Rad Laboratories). Agarose plug was incubated in 275 μl of EPS buffer (0.36 M EDTA pH=8, 0.9% Sarkosyl, 3.6 mg/ml of Proteinase K) at 50 °C for 18 h then washed with TE buffer 3 times for 1 h followed by 3.5 h. Then agarose plug was incubated in 1ml of 0.5M MES buffer (pH=5.5) at 68 °C for 20 min, cooled to 42 °C, and then 1.5 μl of Beta-Agarase (M0392, NEB) was added and incubated for 18 h. Extracted DNA was combed on silianized coverslips (Genomic Vision) using combing machine (Genomic Vision). Combed coverslips was dehydrated at 65 °C for 4 h, denatured in 0.5M NaOH and NaCl for 8 min and dehydrated in a series of ethanol bath for 5min each. CldU or IdU was immunofluorescently detected using an anti-BrdU antibody that recognizes CldU (MCA2060, Bio-Rad Laboratories, 1:50 dilution) or an anti-BrdU antibody that recognizes IdU (347583, BD Biosciences, 1:10 dilution). Combed coverslips were incubated with BrdU antibodies diluted in BlockAid(B10710, Thermo Fisher) at 37 °C for 1h, followed by incubation with Goat anti-mouse IgG Alexa488 (A11029, Thermo Fisher, 1:50 diulution) and Goat anti-Rat IgG Alexa555 (A21434, Thermo Fisher, 1:50 dilution) diluted in BlockAid at 37 °C for 30 min. Images were acquired on a Zeiss AxioObserver Z1, 63 X objective and DNA lengths were measured using ZEN software.

### Re-replication and FACS

Cells were treated with 0.3 μM of MLN4924 for 20 hrs. and fixed with 70 % Ethanol. Cells were suspended in 20 μg/ml of propidium iodide (PI) in PBS with 100 μg/ml RNase A. Fluorescence was measured using a BD LSRFortessa and cell cycle phase was estimated from histograms using Flow Jo.

### SNS-seq and mapping of origins to genome browser

These methods have been described in detail in (31,32). In brief, SNS-seq peaks for each of two biological replicates for a given cell-line were called using MACS2 and peaks found in both replicates (intersection of the two sets of peaks) were considered for further analysis as origins in that cell-line. The median interpeak distance along the reference genome between all peaks was calculated and all peaks closer than this were clusters (initiation zones). To compare the two conditions, we merge the replicate datasets and calculate the number of reads per million (RPM) found in each origin for each cell line. An origins is considered up- or down-regulated if the RPM/origins is 1.5 fold increased / decreased relative to WT cells.

### Statistical analysis

All data analyses were conducted in R 4.1.3 with the ‘tidyvers’(33), ‘Biostrings’(34), ‘BSgenome.Hsapiens.UCSC.hg38’(35), ‘coin’(36), ‘effsiz’(37), ‘ggplot2’(38), ‘rcompanion’(39) packages. If the Shapiro-Wilk test showed that the data were normally distributed, the Welch’s t-test was applied to test for mean differences, and the Hedges’g were calculated to measure the effect sizes. If the data were not normally distributed, or if the number of data points exceeded 5000, the Wilcoxon rank sum test was applied, and the Cliff’s delta was calculated to measure the effect sizes. In the case of multiple comparisons by repeated testing, the Benjamini-Hochberg procedure was applied to obtain adjusted p-values. All statistical tests are two-sided.

### Permutation test

The analysis was performed in R (version 4.1.3), using the ‘tidyvers’(33), ‘regioneR’ (40) and ‘plyranges’(41) packages. RefGene and RepeatMasker data were downloaded from UCSC browser. Data for R-loop, G-quadruplex and MCM subunit ChIP-seq were downloaded from R-loopBase (42), G-quadruplex (43) and ReMap2022 (44), respectively. A permutation test was used to assess the enrichment of origins within the targets. The position of the origins was randomized 1000 times within the genome while retaining canonical chromosomes and avoiding overlaps with masked regions. The number of overlaps between the experimentally observed origins and the targets was calculated. This calculation was also applied to the 1000 randomized set of origins. Statistical significance was assessed by comparing the observed overlap of the experimental data with the distribution of overlap obtained from the randomized data. To quantify the enrichment of origins within the target, fold enrichment (ratio of observed overlap to mean randomized overlap) and z-score were calculated. To evaluate the differences in fold enrichment in the permutation tests (in **Table S2**) between WT origin sets and others, the enrichment of the observed overlap relative to each of the 1000 random overlaps is calculated and then a t-test was examined to see if the differences in the enrichment values of the two sets of origins were significant even after correcting for multiple hypothesis testing (adjusted p<0.05).

### Replication timing

Replication timing data from all 60 samples were downloaded from GSE63428 (45), and Fsu/nimblegen human 2.1M whole genome tiling array data was downloaded from GPL9973. Median values of replication timing for each probe ID were calculated and probe IDs were converted to genomic positions. Genomic positions were lifted over from hg18 to hg38. If the size change was beyond 5%, that locus was discarded. Origin peaks from each sample were intersected to replication timing data. To evaluate similarity of replication timing of origin peaks for each sample, the Kolmogorov–Smirnov test was perfomed and also Wasserstein distance was calculated (46).

### Venn diagram generation

Venn diagrams and a UpSet plot were generated using R (version 4.1.3), with the ‘tidyvers’(33), ‘VennDiagram’(47) and UpSetR (48) packages. The Venn diagram was used to visualize the overlap between origins identified in various samples. The VennDiagram ‘findOverlapsOfPeaks’ function was used to identify overlapping regions, and the minimum number of involved peaks in each overlapping group included in the overlapping counts (’connectedPeaks=“min”).

### Identification of up- or down-regulated peaks in ORCKO cells

Overlap of up-regulated or down-regulated peaks, and peak merging was performed using bedtools v2.29.0. The origins up- or down-regulated in the knockout (relative to WT) were intersected with each other. If such peaks from all knockout cell lines overlapped with each other, they were merged. CommonKO_up refers to the up-regulated peak, and CommonKO_down refers to the down-regulated peak.

## RESULTS

### Survival of ORC knockout cells without re-expressing ORC

If a cell survives at a disadvantage because of the absence of ORC subunits, one possibility is that the protein (or a truncated, functional protein) is somehow re-expressed to allow the cells to survive. This was not the case for the ORC knockout cells. The levels of ORC1, ORC2 and ORC5 in the *ORC1Δ*, *ORC2Δ* and *ORC5Δ* cells were still undetectable on Western blots of cell lysates even though these cell lines were derived years ago, and each of them have been cumulatively passaged for several months (**Figure 1A)**. The deleted ORC subunits remained undetectable on the chromatin fraction in both low passage cells and high passage cells (**Figure 1B)**. ORC1 association with the chromatin is decreased when *ORC2* or *ORC5* are deleted, but that could be partly due to destabilization of the protein in those cells (levels decreased in input cell lysates in **Figure 1A**). ORC2 association with the chromatin is not affected by *ORC1* or *ORC5* deletion. ORC5 association with the chromatins decreased not only when *ORC5* is deleted, but also decreased due to destabilization (decreased in the input cell lysate) when *ORC2* is deleted.

**Figure 1.**
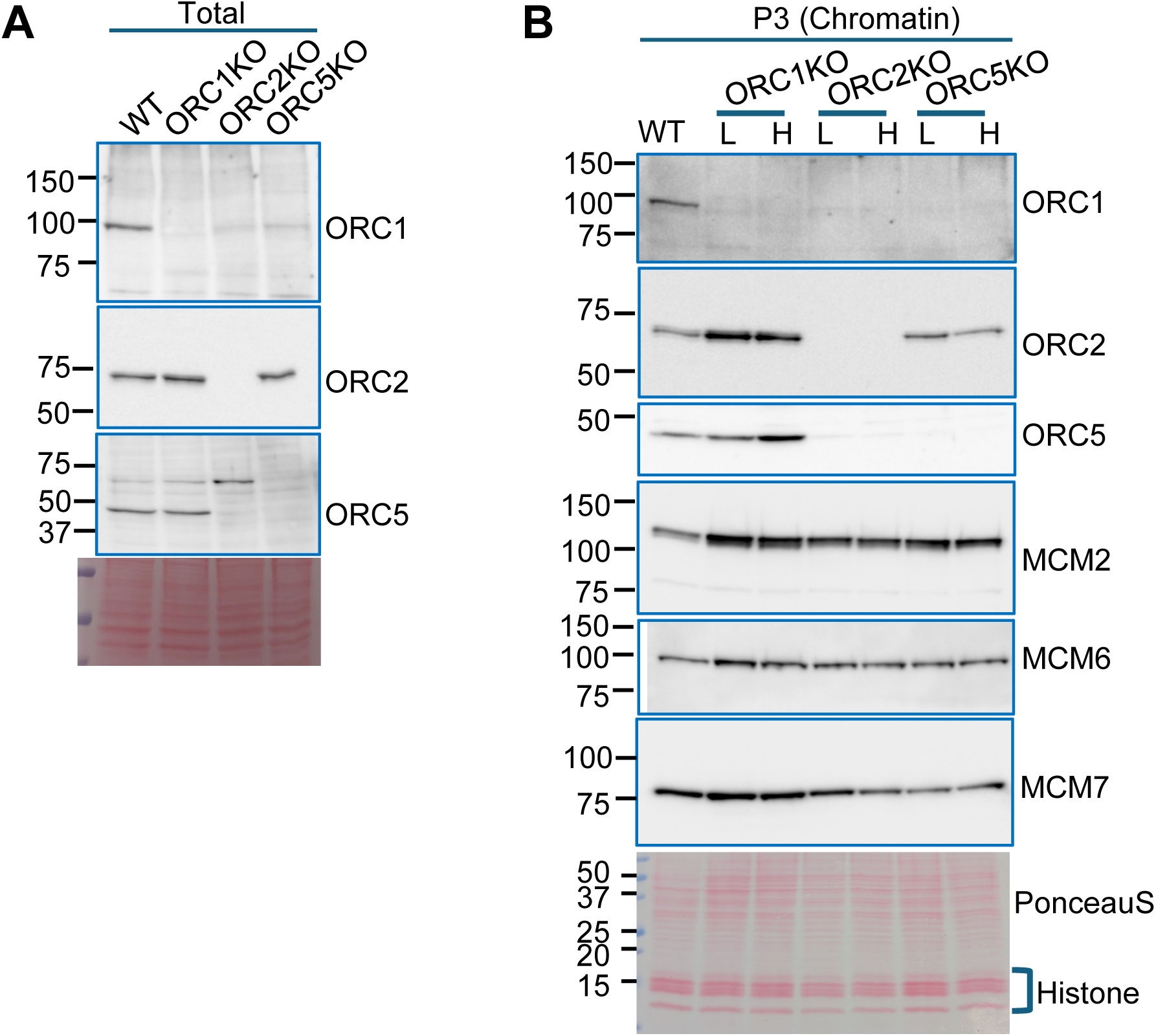
Chromatin loading of MCM2-7 in ORC KO cell lines. A) Immunoblot of total protein in the ORC KO cell line. B) Immunoblot of chromatin-associated proteins in the ORC KO cell lines. L: Low passage, H: High passage

We had reported that MCM2-7 loading on chromatin was decreased compared to WT cells in the first report of *ORC1Δ* and *ORC2Δ* cells, even though it was sufficient to fire a normal number of origins (24). In a later paper we noticed that the MCM2-7 loading on chromatin did not decrease in the *ORC5Δ* and *ORC2Δ+ ORC5Δ* cells (23). Now we have carefully compared the MCM2-7 loading from low (<1 month) and high (>3 month) passage cells of all three KO cells and find that MCM2-7 is always associated with the chromatin at levels in the *ORC1Δ* or *ORC2Δ* that are comparable to that in WT cells or the *ORC5Δ* cells (**Figure 1B**). We do not know why *ORC1* or *ORC2* deletion appeared to have a bigger effect on MCM2-7 loading when we first examined this question (24), but this does not change the conclusions of that paper: enough MCM2-7 was being loaded to license sufficient origins for normal S phase progression and for cell proliferation. We are still left with the following questions regarding MCM2-7 loading in the absence of the six-subunit ORC: Is the MCM2-7 being loaded at specific sites to license specific origins? Is it being loaded at new sites in the absence of ORC subunits? Is an excess of *functional* MCM2-7 being loaded in each cell-cycle as it does in the presence of ORC? Can functional MCM2-7 be loaded *multiple* times in the same S phase despite the absence of ORC?

### The majority of the origins map to WT sites in the ORC knockout cells

SNS-seq is a well- accepted method to identify origins of replication in a population of cells (49–52). All four cell lines yielded robust SNS-seq peaks (**Figure 2A**). Origin selection in mammalian cells is can sometimes vary in different cell lines, although significant overlap is detectable (53). Based on our analysis of the published literature, we believe that Core origins described from 19 human cell samples by Akerman et al. (54,55) constitute the most reliable set of origins to use as a benchmark. Although the Core origins were not derived from HCT116 colon cancer cells, about 46% of the origins in Wild type HCT116 cells overlapped with the core origins (**Figure 2B**). Core origins are origins from which 80% of replication initiation events occur in various human cell types analyzed, such as pluripotent cells, primary cells, differentiated cells, and immortalized cells. We wondered whether the origins mapped in WT HCT116 (colon cancer epithelial cells) will overlap more with origins present in epithelial cells. The overlap of HCT116 origins with primary or “p53KD+RAS immortalized” or “p53KD+WNT immortalized” breast epithelial cells was lower (27%, 34% and 29%, respectively) than that with the Core origins.

**Figure 2.**
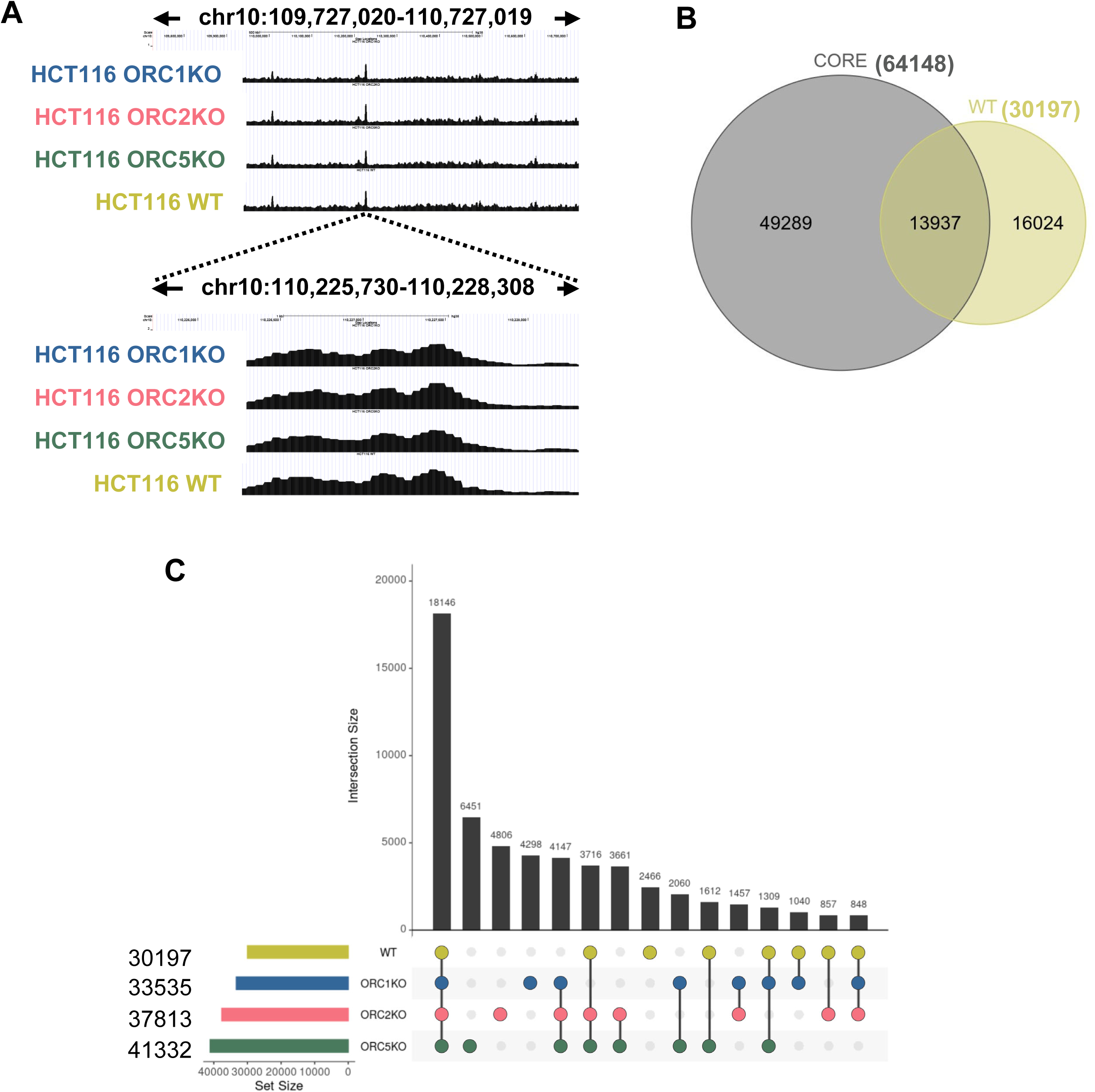
A) An example of a typical origin peak seen in WT and ORC subunit knockout cells as seen with the UCSC browser. Each black colored region indicates origins from two replicates of each cell line. B) C) Overlap of origin peaks in WT and ORC subunit knockout cells. Numbers in parentheses indicate the total number of peaks in each cell lines. Because multiple peaks from one group may overlap with a single peak from the other, the overlapping peak number is the minimum number of overlapping peaks from either of the two overlapping groups. (B) Overlap between CORE origins and origins from WT cells. (C) Overlap of origins derived from WT and ORC subunit knockout cells.

Nearly 60% of the origins mapped in WT cells were detected in all three ORC knockout cells (**Figure 2C**). It was equally interesting that over 90% of the origins mapped in WT cells were seen in one or more of the ORC knockout cells. This degree of overlap in origins between the four cells suggests that cells are mainly using the same set of origins in the presence or absence of ORC or of some of the subunits of ORC on chromatin.

We have reported that loss of one subunit of ORC decreases the loading of the other subunits in a very specific manner (23,24). The *ORC2Δ* cells had a great decrease of 4 subunits (ORC2, ORC3, ORC4 and ORC5) and a moderate decrease of 1 subunit (ORC1) on chromatin. The *ORC5Δ* cells had a great decrease of 2 subunits (ORC4 and ORC5) on chromatin. The *ORC1Δ* cells showed a decrease of only 1 subunit (ORC1) on chromatin. 21,343 origins (71% of WT origins) were common between WT and *ORC1Δ* cells. 23,567 origins (78% of WT origins) were common between WT and *ORC2Δ* cells. 24,783 origins (82% of WT origins) were common between WT and *ORC5Δ* cells. Thus, the different subcomplexes (varying from 2 to 5 subunits) that remained on the chromatin in the different mutant cells did not select very different subsets of origins.

### Subsets of origins are commonly up-regulated or down-regulated after knockout of individual ORC subunits

We next focused on origins that were selectively upregulated (ORC KO-up) or downregulated (ORC KO-down) in the knockout cells, relative to the WT cells (**Figure 3A**). The Venn diagrams show that 4000 origins were commonly up-regulated in the three different knockout cells, while 2500 origins were commonly down-regulated in all three knockout cells (**Figure 3B**).

**Figure 3.**
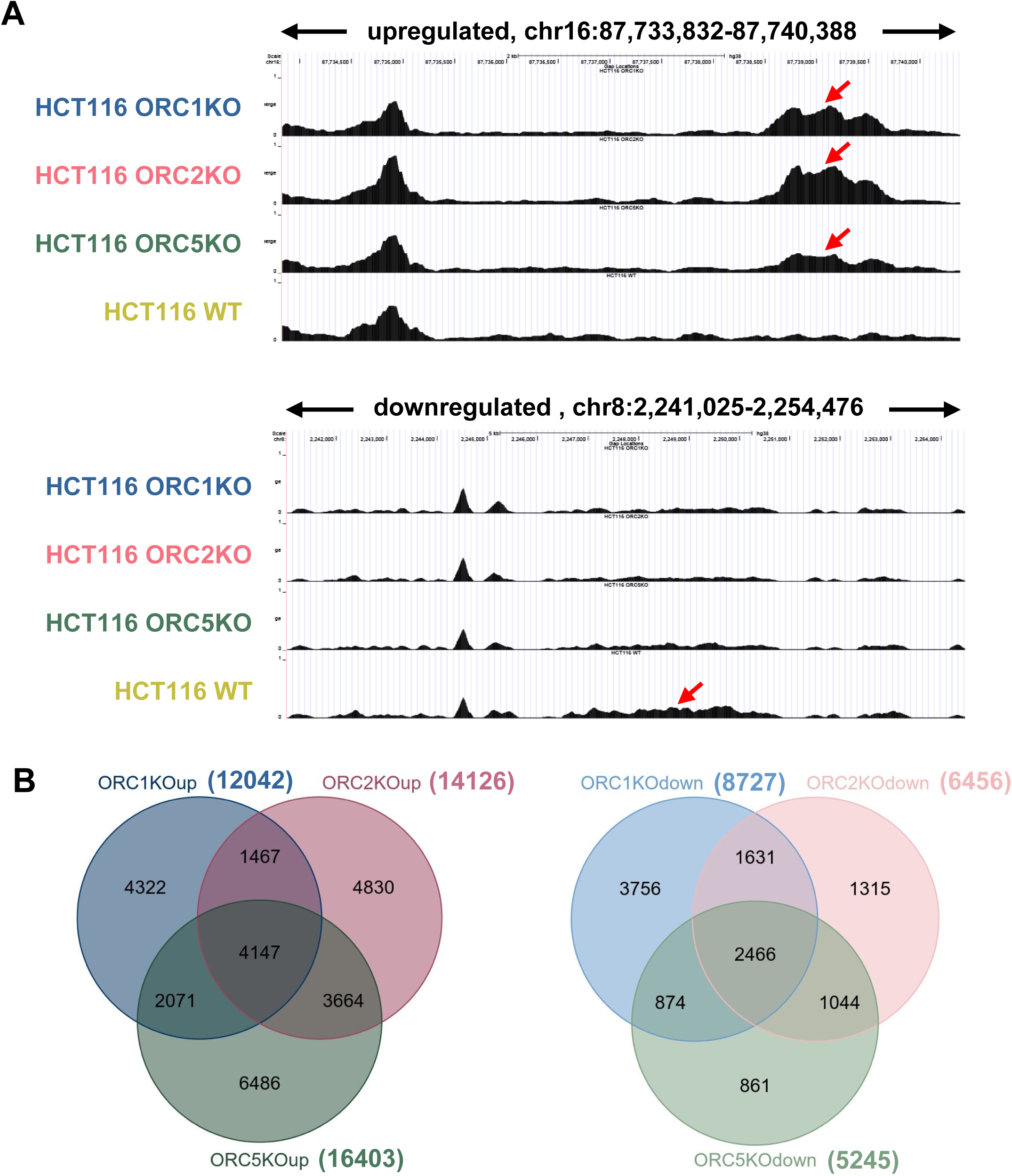
A) Examples of a typical up-regulated or down-regulated origins seen in ORC subunit knockout cells as seen with the UCSC browser. Each black colored region indicates origins from two replicates of each cell line. The red arrows indicate differential origins. B) Venn Diagram of origins that were up-regulated in ORC subunit knockout cells (compared to WT cells) to show overlap between the knockout cells (left). A similar analysis was done for origins that were down-regulated in the knockout cells (right). Rest as in Fig. 2.

### Properties of the origin sets

The DNA near origins has been reported to have high nucleotide distribution skew (56). To compare the properties of the origins in WT and ORC knockout cells, we examined whether they were significantly different for SNS-seq peak width (origin length) as well as origin GC content, TA skew, and GC skew by Welch’s t test or Wilcoxon rank sum test (see Materials and Methods). GC skew measures the over-abundance (positive value) or under-abundance (negative value) of G relative to C on one strand of the DNA, while absolute GC skew removes the negative sign and indicates whether there is an imbalance of G relative to C. TA skew measures similar imbalance for T relative to A. The width (length) of the origin peaks (**Figure 4A**), GC content (**Figure 4B**) and TA skew (**Figure 4C**) were not different between origins in WT cells and the Core origins (48,49). These properties were also similar between origins in WT and ORC knockout cells, even though some of the differences were statistically significant (**Supplemental Table S1**). However, the GC skew (**Figure 4D**) was much higher (effect size ≥0.554) in the experimental HCT116 cell lines, both WT and the knockouts, compared to the Core origins. **Figure 2B** shows that around half of the origins present in WT cells are specific to HCT116 cells. Perhaps such HCT116 specific origins are responsible for the difference in GC skew between CORE and WT, but there was not much of a difference between all the origins in the knockout and WT cells (**Supplemental Table S1**, lines 3-6).

**Figure 4.**
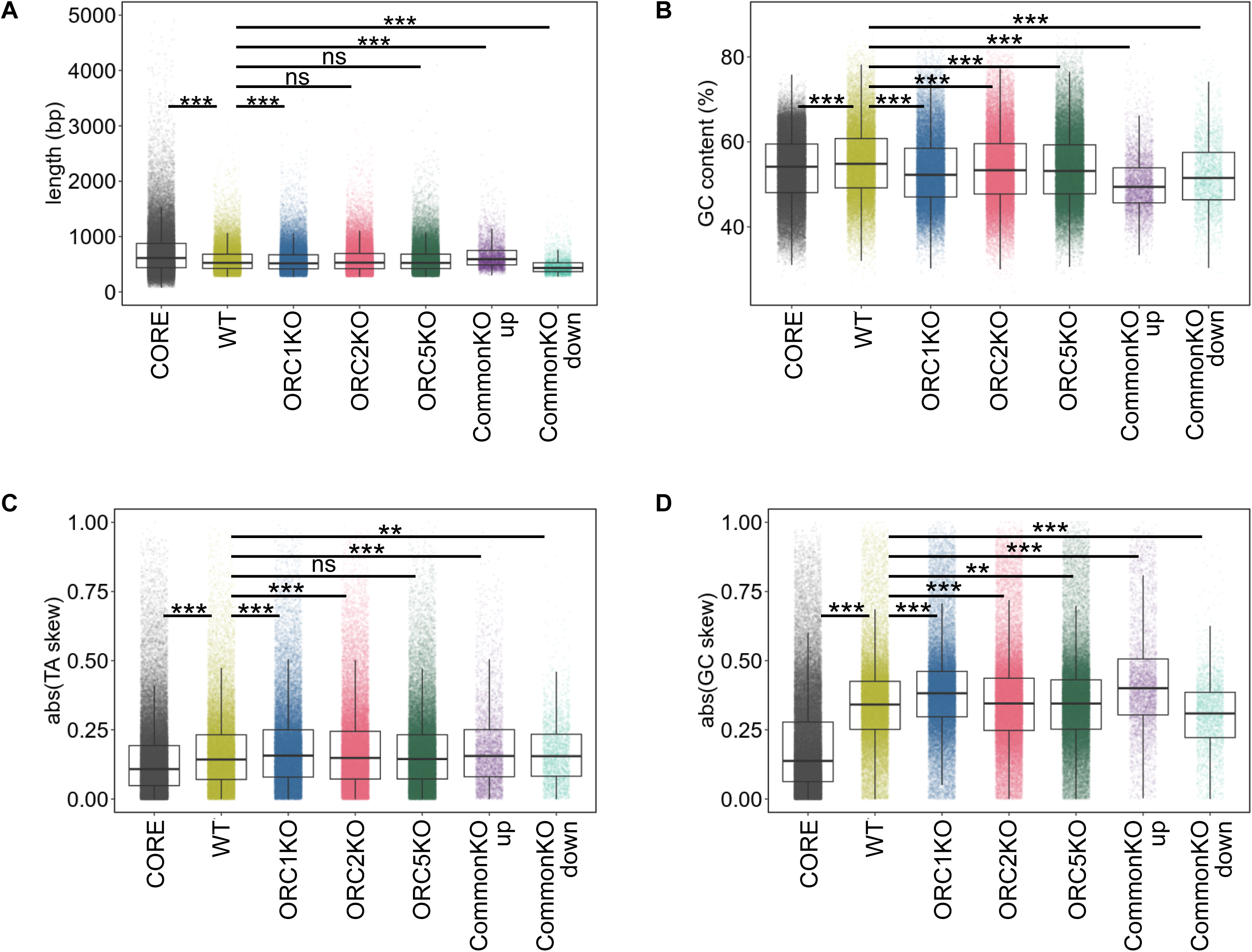
Distribution of (A) length, (B) GC content, (C) TA skew and (D) GC skew for origins in the WT, knockout cells, differentially regulated origins in knockout cells (compared to origins in WT cells) and CORE origins. Box plot shows median and interquartile range. For the TA skew and GC skew, the values were converted to absolute values, (abs(TA skew), abs(GC skew)). P values were determined by two-sided Wilcoxon rank sum test with the Benjamini–Hochberg procedure (FDR correction). * adjusted p-value < 0.05, ** adjusted p-value < 0.01, *** adjusted p-value < 0.001, ns means not significant. CommonKO_up refers to the up-regulated peaks in all knockout cell lines, and CommonKO_down refers to the down-regulated peaks in all knockout cell lines.

In contrast, when we analyzed the origins that were up-regulated or down-regulated in *all* three knockout cells (the overlap group indicated in **Figure 3B**), some differences emerged (**Figure 4 and Supplemental Table S1**). We did the Wilcoxon rank sum tests to obtain p-values and also calculated Cliff’s Delta to measure the dissimilarity of the two datasets (e.g. parameter of origins in the two cell lines being compared) (**Supplemental Table S1**). The differences we cite below are statistically significant and have a measurable dissimilarity (effect size). While origins in any of the KO cells are very similar in length compared to WT cells (negligible Cliff’s Delta), the CommonKO_up origins are longer and CommonKO_down are shorter than the origins in WT cells. Similarly, the CommonKO_up and CommonKO_down origins have lower GC content than the other sets of origins. The GC skew in the CommonKO_up was distinctly higher than that in WT origins (even though all origins in the *ORC2Δ* and *ORC5Δ* cells are not different from WT origins). Conversely the GC_skew in the CommonKO_down set was distinctly lower than in WT origins. The result suggest that the new origins activated in common by all three ORC subunit deletions (CommonKO_up) appear to be slightly but significantly longer, with lower GC content and higher GC skew than WT origins or the pool of all origins in the KO cells.

### Origins in WT and KO cells are enriched in MCM binding sites

The MCM2-7 complex is loaded on chromatin at DNA replication origins (57). We examined whether the MCM2-7 complex is enriched on DNA replication origins in ORC subunit-deficient cells by permutation tests. Binding sites of MCM2, 3, 5, and 7 subunits determined in ORC proficient cells (44) were enriched on the origins in both WT and ORC knockout cells (**Figure 5A**). The significance of any difference in the enrichment of a MCM binding site between the origin sets was determined following the permutation test as described in the methods (**Supplementary Table S2**). All the sets were statistically significantly different from the WT origin set, but the effect sizes (the magnitude of the difference in enrichment) were mostly negligible. Relative to all other origin sets, the origins up-regulated in all knockout cells (CommonKO_up) were most enriched for MCM2-7 binding sites, while the origins down-regulated in all knockout cells (CommonKO_down) were least enriched in MCM binding sites (**Figure 5A, Supplementary Table S2**). Thus, even in the absence of a functional ORC subunit, the active origins are enriched in sites where the MCM2-7 complex is known to be bound to the DNA in WT cells, suggesting that loss of ORC does not result in a large redistribution of MCM binding sites. The higher enrichment of MCM binding sites at origins that are activated in the absence of ORC suggests that the high probability of MCM binding to these sites is determined independent of ORC, and this higher probability becomes significant in firing new origins when ORC is absent.

**Figure 5.**
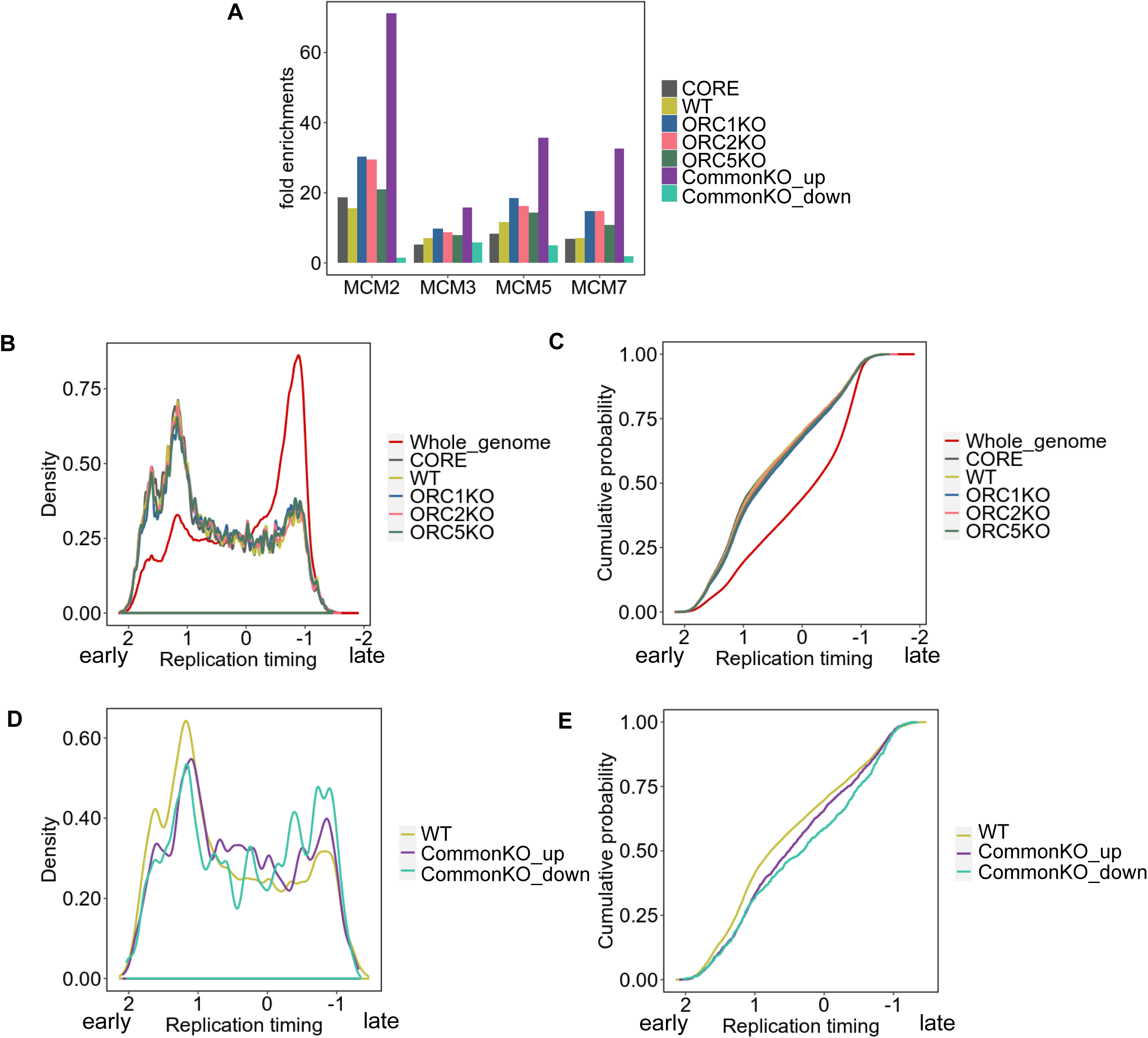
Comparison of origins in WT and ORC subunit knockout cells with MCM loading and time of replication in S phase. (A) Permutation test results of colocalization between various origin peaks and MCM subunits (reference). 1,000 randomized region sets were obtained for each origin set and intersected with peaks of each MCM subunit.□ The mean of the overlap seen with the 1000 random sets is the expected overlap.□ The Y-axis indicates fold enrichment of the experimentally determined overlap with this expected overlap. (B-E) Distribution of origins as a function of replication timing (see Materials and Methods).

### Replication timing of origins in knockout cells

SNS-seq studies suggested that DNA replication origins tend to be enriched in early replication regions. We examined whether this holds true in the absence of functional ORC subunits by plotting the distribution of the origins against the time of replication of the co-located part of the genome (**Figure 5B-D**). The distribution curves in Figures 5B,C were compared by two metrics to quantify whether the curves were different from each other. KS-distance, defined as the maximum vertical distance between two empirical cumulative distribution functions, has several limitations, including high sensitivity to differences between the modes of the two distributions but insensitivity to their tails. In contrast, the Wasserstein distance (WD) is a function of the horizontal distance between the observations, and is a better measure of the effect size. As expected, origins in WT cells were enriched in the early replication regions compared to the distribution of all genomic loci (**Figure 5B, C; Supplementary Table S3**). However, the timing of replication of origins in the knockout cells was not very different than in WT cells (WD is small). On the other hand, the distribution of replication timing of origins that were up- or down-regulated in all the knockout cells was slightly different from that of WT cells. Those origins were more enriched in the mid and late replication parts of the genome (**Figure 5D, E, Supplementary table S3**).

### Overlap of origins with known genomic annotations

It is known that Core origins are enriched in promoters of genes (2kb upstream of genes, exons, 5’UTRs and CpG islands (55). We examined whether this was also true for the origins in WT and ORC knockout cells by permutation tests. As already reported, the Core origins were found to be more enriched in these four areas than expected from the random distribution (**Figure 6A**). These properties are preserved in all four experimentally determined origin sets (WT, and the three knockout cells) and also in the origins that are downregulated in all three *ORCΔ* cells. However, a surprising exception is seen for the origins that are upregulated in all the *ORCΔ* cell lines, where there is a much less enrichment over promoters, 5’UTRs, exons or CpG islands (**Figure 6A, Supplementary Table S2**).

**Figure 6.**
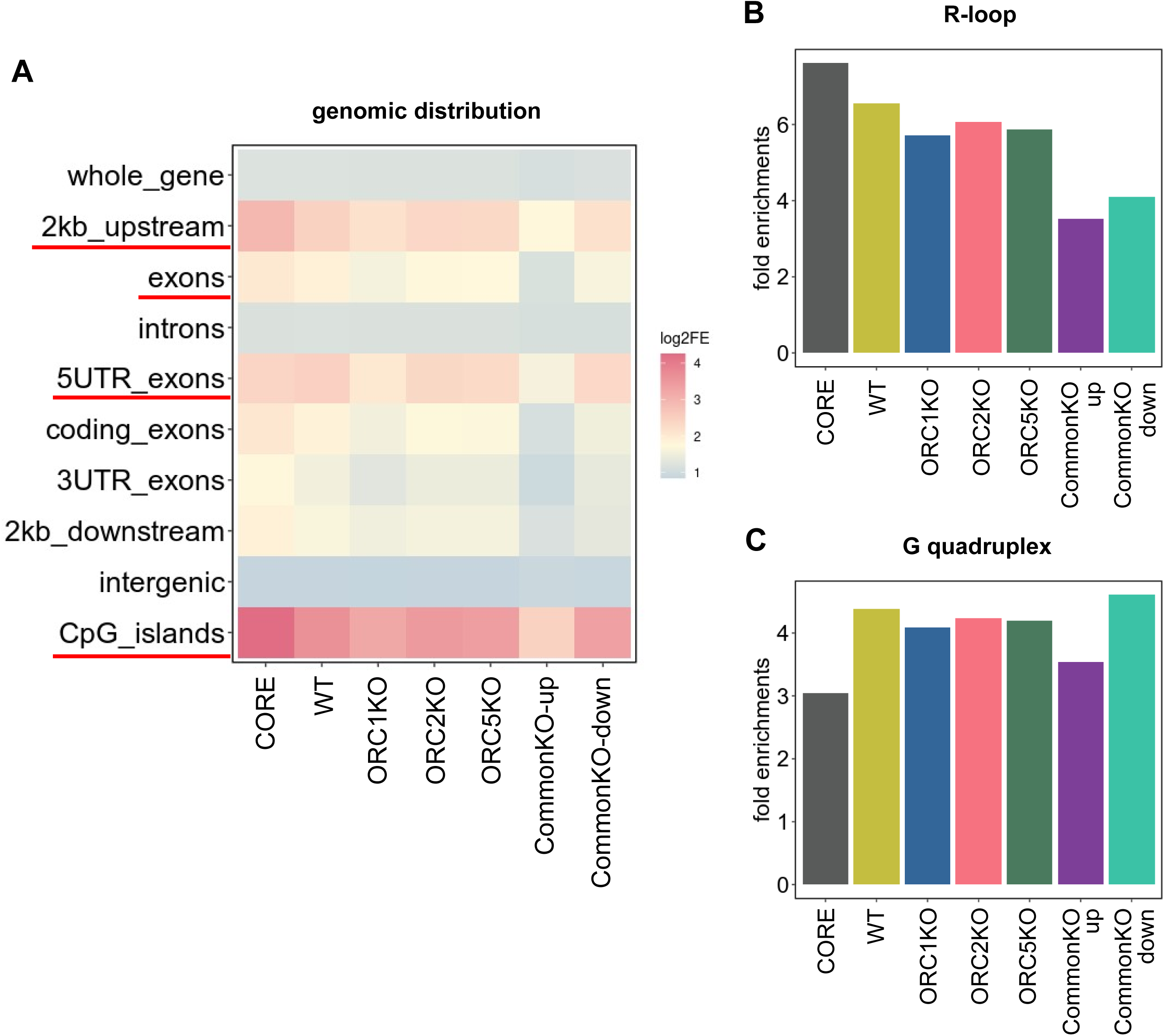
Permutation test to determine whether the colocalizations indicated are enriched relative to random expectation.□ Colocalization of sets of origins and (A) indicated genomic sites, (B) Level 5 R-loops from (42) and (C) G quadruplexes (43) (G4H1.52). The rest as in Figure 5A.

As reported before, origins are also enriched over random expectations in areas that form R-loops and G-quadruplexes (**Figure 6B, C**) (58,59). This was true for all the origins in the WT and knockout cell lines. The enrichment is still seen but is lower for the origins that are up- regulated in all of the ORC knockout cells (**Figure 6C, Supplementary Table S2**), and this lower enrichment at G-quadruplexes is likely due to the lack of enrichment of these origins near genes and promoter. These results suggest that origins in HCT116 WT cells, and most origins seen in ORC knockout cells, with some minor differences, have properties similar to CORE origins. There is, however, a consistent difference in CommonKO_up origins, which are less-enriched for promoters, 5’UTRs, exons or CpG islands, R loops or G quadruplexes than all origins in WT or KO cells. The CommonKO_down origins were similar to most of the origins in WT and the KO cells except that they were less enriched in R loops.

### Overlap of origins with (ATCC)n repeat elements

Repeat elements are often enriched near bacterial and eukaryotic origins (58,60). We tested whether any of the classes of origins we have identified are significantly different from the general class of origins in terms of their overlap with repetitive elements by permutation tests. The full set of origins we identified in the WT and knockout cells were enriched relative to Core Origins in low complexity sequences and in simple repeats (**Figure 7A, B**). The differences in enrichment of repetitive elements with all the origins in the knockout cells compared to that with the origins in the parental WT cells were statistically significant but very small (**Supplementary Table S2**). The exception was when the overlap of origins with (ATCC)n motifs (and permutations or complementary sequences) were compared, where we find that the CommonKO_up were specifically highly enriched, while the CommonKO_down were specifically dis-enriched in (ATCC)n motifs (**Figure 7C, D**). The enrichment of CommonKO_up with (ATCC)n motifs could be significant because a negative control, (TACC)n motif, was not more enriched in the KO-up origins compared to the other sets of origins in WT or KO cells (**Supplementary Figure S1**).

**Figure 7.**
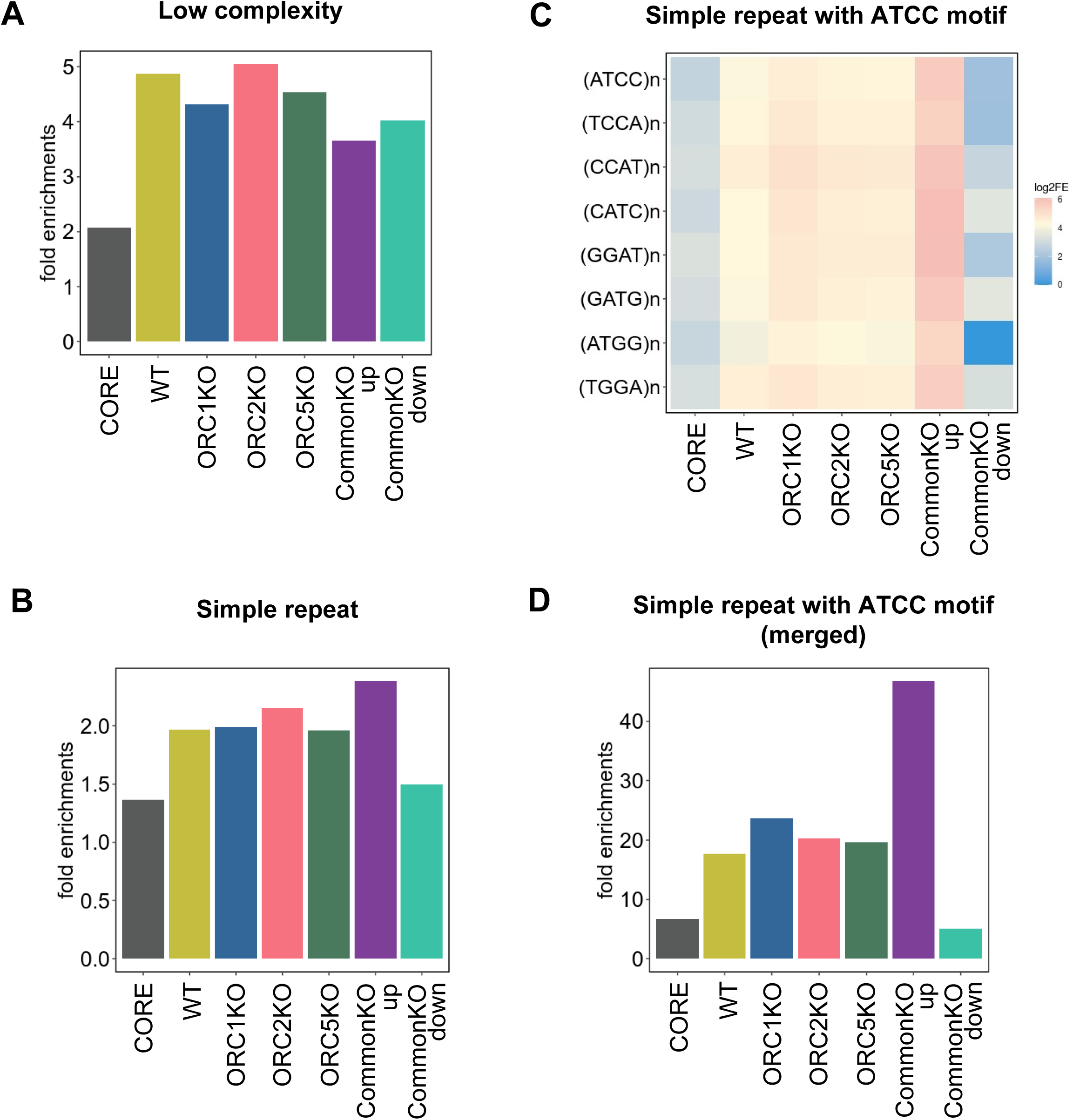
Permutation test results of colocalization between various origin peaks and (A) low complexity regions, (B) simple repeat regions, (C) each simple repeat regions including various type of ATCC motif and (D) merged simple repeat regions with ATCC motif. The rest as in Figure 5A.

### Measurement of functional MCM2-7 loading in the ORC knockout cells

A flow- cytometry based approach to measure the MCM2 loading on DNA per cell was developed by Jean Cook and collaborators (30). We used this approach to independently confirm that the same maximal level of MCM2 loading on DNA in G1 was seen in the *ORC1Δ*, *ORC2Δ* and *ORC5Δ* cells as in the WT cells (**Figure 8A**).The FACS plots also show that the normal progressive loss of MCM2 from the chromatin during S phase was seen in the mutant cells in a manner similar to the WT cells. Finally, this FACS approach allows us to quantify the relative MCM loading rate in WT and ORCΔ cells using ergodic rate analysis (30,61) (**Figure 8B**). There was no significant difference in MCM loading rate between WT and ORC1Δ or ORC2Δ cells and ORC5Δ cells were about twice as fast as WT cells.

**Figure 8.**
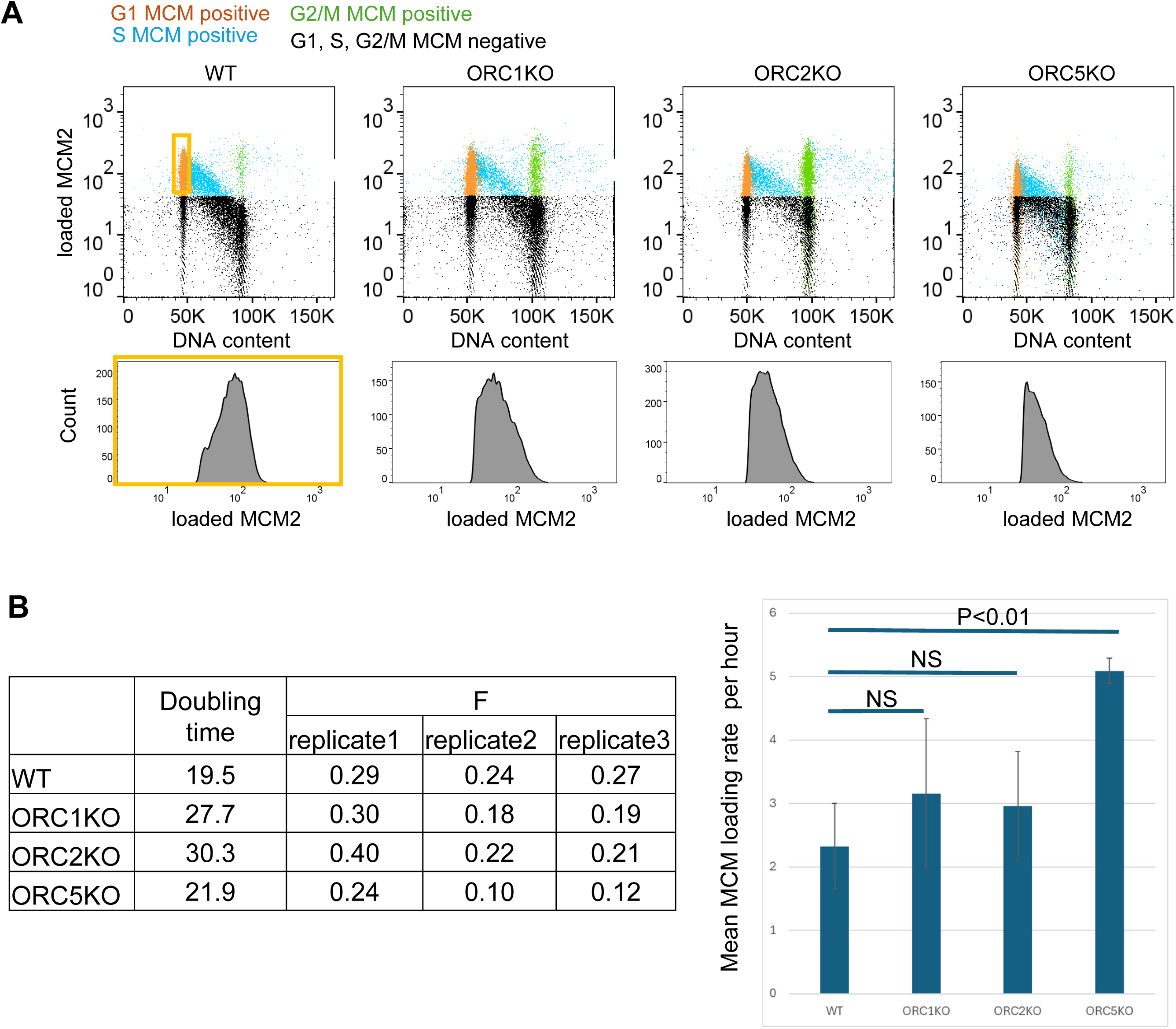
A) Top: Flow cytometry plots of MCM2 loading on DNA (Y axis) against DNA content as measured by DAPI (X-axis). The yellow window marked as G1, MCM positive, contains G1 cells that have loaded MCM2 on DNA. Bottom: Distribution of MCM2 staining intensity and cell count from the G1 populations of indicated cell lines. The maximum X-axis value for each curve is a measure of the maximum amount of MCM2 loaded on chromatin at the end of G1 in that cell line. Profiles are representative of two independent experiments. B) Left: Parameters used to calculate Right: Calculated mean MCM loading rate per hour by ergodic rate analysis (n = 3) from two independent experiment, unpaired two tailed t-test. Data are shown as mean□±□s.d.

A 3-10 fold excess of MCM2-7 double hexamers is loaded on chromosomes at the end of G1 than is necessary to fire the ∼50,000 origins that are required to completely replicate the genome in each S phase. Normally, the excess loaded MCM2-7 hexamers are passively displaced from chromatin by replication forks originating in adjoining DNA and replicating through the unfired MCM2-7 hexamers. However, the excess MCM2-7 double hexamers provides dormant origins that become important under conditions of replication stress when replication forks stall before traversing their normal distance. The dormant origins in the unreplicated parts of the genome now get activated and this is necessary to replicate the entire genome despite the premature stalling of replication forks (6,7,62).

To measure this origin provided by the dormant but licensed origins, we put the WT and ORC deletion cells under replication stress by exposing them to low doses of hydroxyurea and following their proliferation by clonogenic assays. The knockout cells were not more sensitive to hydroxyurea compared to WT cells, suggesting that there was a normal complement of dormant origins in the knockout cells (**Figure 9A**). Because normally unfired origins are now initiating DNA replication, the firing of dormant origins decreases the inter-origin distance, as measured by molecular combing. Consistent with this, the inter-origin distance decreased upon exposure of WT cells to HU, and a similar decrease was seen in the cells where *ORC2* or *ORC5* were knocked out (**Figure 9B**). Therefore, excess functional MCM2-7 was still loaded (compared to the minimum required to license ∼50,000 origins), and the dormant origin function was intact despite the absence of ORC.

**Figure 9.**
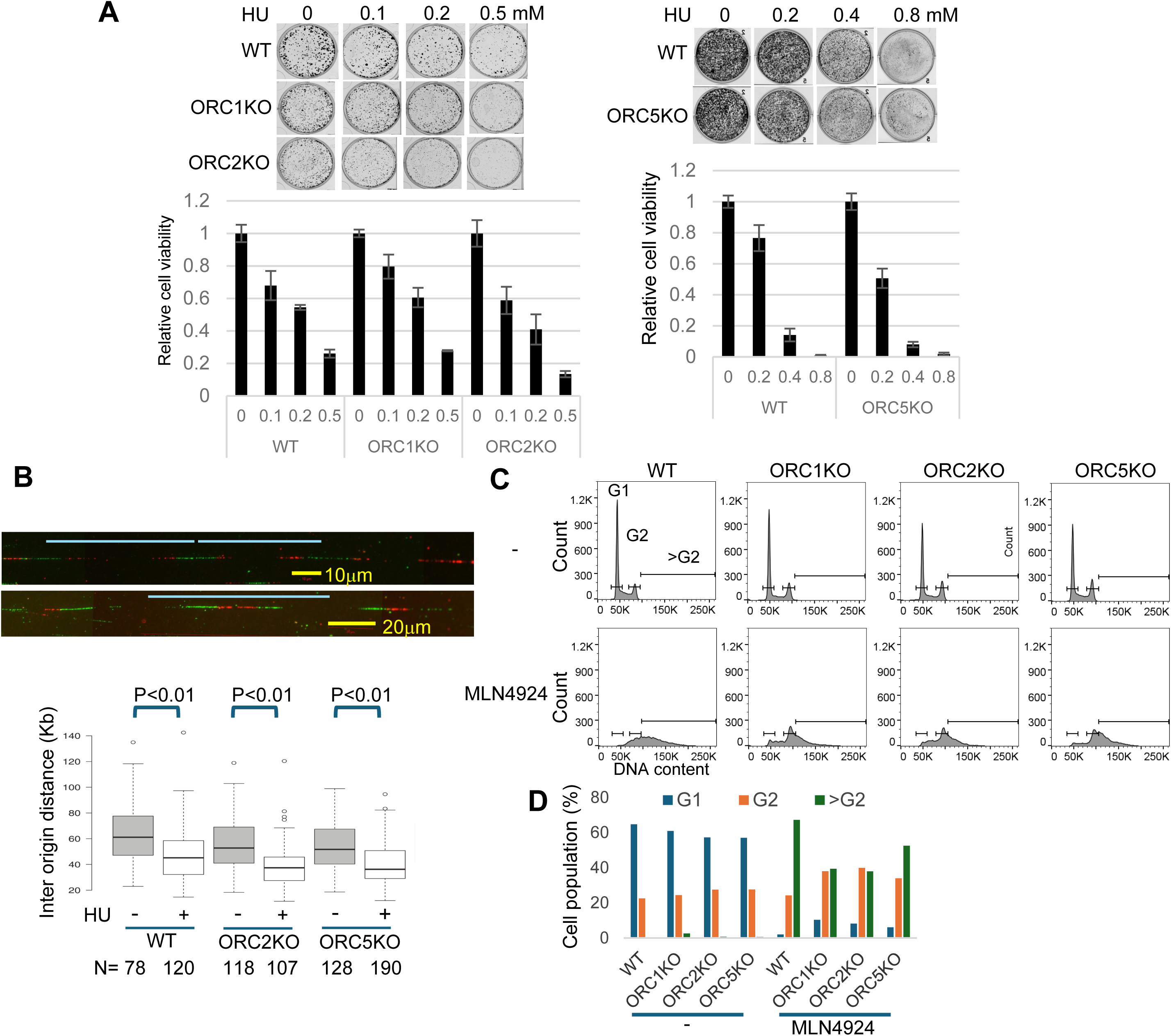
DNA replication in ORC KO cell lines. A) Clonogenic assay of Hydroxyurea treated ORC KO cell lines. Cells were treated with indicated amount of Hydroxyurea for 7 days and colonies detected by Crystal Violet staining (Top). Crystal violet-stained colony density was measured. Data was normalized to the colony density of Day 1 (Bottom). Analysis (n = 3) from single experiment, unpaired two tailed t-test. Data are shown as mean□±□s.d. B) DNA molecular combing of cells treated with or without Hydroxyurea for 4 hrs. (Top) representative image. Blue line indicates inter origin distance. (Bottom) Box and whiskers plot for inter-origin distance of indicated clones of cells. Two-sided Wilcoxon rank sum test for two samples N=number of DNA fiber from single experiment. P: Statistical significance of any difference between HU untreated and HU treated cell. Data are shown as median□±□s.d. C) FACS profile of propidium iodide-stained cell treated with MLN4924 for 20 hrs. Profiles are representative of two independent experiments. D) The percentage of cells in G1, G2, >G2 in (C) is plotted. Profiles are representative of two independent experiments.

We and others have also reported that an artificial increase in the amount of replication initiation factor, Cdt1, relative to an inhibitor of Cdt1, geminin, leads to repeated licensing and firing of origins on already replicated parts of the genome in the same S phase, resulting in re-replication of the DNA (63,64). This re-replication leads to the production of cells with greater than G2 DNA content and of course, is dependent on the loading of functional MCM2-7 in S phase on parts of the genome that have already replicated once and lost the MCM2-7 that was loaded in G1. Treating cells with MLN4924 inhibits the degradation of Cdt1, increases the amount of Cdt1 relative to geminin and leads to re-replication (65). Re-replication upon addition of MLN4924, as seen by the appearance of cells with >G2 DNA content, was seen in the *ORC* knockout cells (**Figure 9C**), consistent with the suggestion that functional MCM2-7 can be loaded in these cells even in the absence of ORC. However, there may be a limit to the number of times MCM2-7 can be re-loaded in the knockout cells, accounting for the 30-40% decrease in the number of cells with >G2 DNA content (**Figure 9D**)

## DISCUSSION

There are two important questions we set out to answer. Given the obvious importance of ORC in binding DNA and loading MCM2-7 double hexamers in the vicinity, the first question was whether the cells without an ORC hexamer still choose specific sites as origins and if they do, whether those origins have properties different from the origins chosen in the presence of ORC. The second question examined the robustness of the chromatin loading of functional MCM2-7 hexamers in cells without the six-subunit ORC. While MCM2-7 is clearly associated with the chromatin, and sufficient for executing enough DNA replication for cell survival, is there still an excess of functional MCM2-7 hexamers to license the dormant origins that become important for cells to complete replication and survive in the face of replication stress. Furthermore, can functional MCM2-7 hexamers be loaded repeatedly on the same stretch of DNA to facilitate re-replication in the ORC deficient cells?

### Presence of a trace, yet functional amount of ORC in the ORC knockout cell

In a very important paper, Chou et al., reported that our *ORC2Δ* cells contain a trace amount of a protein smaller than ORC2 that co-immunoprecipitates with ORC3 and reacts to their anti-ORC2 antibody (25). They suggested that the *ORC2Δ* cells may produce a trace amount of an N terminally deleted ORC2 protein due to initiation from an internal methionine. They could not detect any ORC1 in the *ORC1Δ* cells. The *ORC5Δ* cells were designed such that after the deletion of the initiator methionine, the next methionine is downstream from the bulk of the AAA+ domain, so that even if a truncated protein was produced from internal initiation at the second methionine, it would miss a major functional domain of ORC5. Thus, there is no controversy over the absence of ORC in the *ORC1Δ* and *ORC5Δ* cells.

Turning to the *ORC2Δ* cells, even if the protein detected in (25) is functional, we quantitated the published Western of the cell lysates (Figure 8, Supplement 1b, from that paper) to estimate how much of the truncated protein was present per cell. The intensity of the signal from the truncated protein in the *ORC2Δ* cells was 0.23% of that of full length ORC2 in WT cells (and this is an overestimate, because the signal for the full length ORC2 protein is highly over- exposed). Quantitative Westerns in (24) estimated that WT cells contain ∼150,000 molecules of ORC2, suggesting that the truncated ORC2 detected by Chou et al. exists at 345 molecules per cells. The inter-origin distance in the *ORC2Δ* cells is ∼40 kb (**Figure 9**, in hydroxyurea), suggesting that ∼150,000 origins are licensed on the 6x10^9^ base-pair genome of a diploid cancer cell. It is difficult to explain how 345 molecules of ORC2 could load ∼150,000 MCM2-7 double-hexamers per cell, given that the *in vitro* MCM2-7 loading experiments (66,67) reveal no evidence of ORC re-cycling in the loading reaction (John Diffley and Franziska Bleichert, personal communication).

Under our best conditions, we do not detect the truncated ORC2 protein in cell lysates or in ORC3 immunoprecipitates of the *ORC2Δ* cells with either the antibody used by Chou et al, or a commercial antibody. We also do not see any ORC2 related bands in that size-range that are knocked down by si-ORC2 (**Supplementary Figure S2).** We acknowledge that such differences can rise from differences in the sensitivity of Western blot protocols, but the result highlights that the protein under consideration is present at very low levels.

### Essentiality of *ORC* genes, particularly *ORC2* gene

Chou et al. show that multiple sgRNAs against ORC subunits are dis-enriched in CRISPR screens on WT cells (25). Furthermore, in cells subsisting on ORC2 fused to an auxin-inducible-degron, rapid degradation of ORC2 diminishes (but does not abolish) MCM2-7 loading on chromatin in WT cells. This is the best evidence showing that ORC2 subunit is essential for cell viability and for MCM2-7 loading in WT cells but does not rule out the possibility that a small fraction of cells can be selected for that can survive long term and load MCM2-7 without ORC2. A more difficult result to explain is why sgRNAs against *ORC2* were dis-enriched even in the *ORC2Δ* cells, suggesting that the *ORC2* gene is still essential for proliferation in the *ORC2Δ* cells. Given that we see proliferation and normal MCM2-7 loading in the same *ORC2Δ* cells despite the complete (or near complete) depletion of ORC2 protein, three possibilities remain: (a) a trace amount of ORC2 protein can be recycled and load many fold excess of MCM2-7 on the chromatin *in vivo*, (b) origins on a few selected parts of the genome, but not most of the origins in the cell, are critically dependent on ORC for loading MCM2-7 or (b) a trace amount of ORC2 protein is required for some other function in the cell that is different from MCM2-7 loading.

The Dependency map (DepMap) (68) measures the depletion of the targeting shRNAs or sgRNAs from the starting library after 40 days of passage of the targeted cells. While shRNAs or sgRNAs for essential genes are depleted nearly completely, giving a Dependency index near 1, shRNAs or sgRNAs that slow cell proliferation, or initially slow cell proliferation but then allow the cells to adapt, will be depleted to an intermediate level, yielding a smaller Dependency index. Keeping this in mind, the Dependency index of several of the ORC subunit genes in cells vary among each other, something not expected if the entire six-subunit ORC complex was essential. In addition, the Dependency index of several of the ORC subunit genes are much lower than 1 in HCT116 colon cancer cells. For HCT116, the Dependency index of *CDC6* and *ORC6* (which we have been unable to target) are 0.988, and 0.971, respectively, while that of *ORC2*, *ORC3*, *ORC4* and *ORC5* are 0.321, 0.133, 0.471 and 0.390, respectively. Our ability to generate viable cells after targeting *ORC1* is more surprising because the dependency index of *ORC1* is 0.961. However, it was reassuring that Chou et al. (25) examined our *ORC1* KO cells and did not find evidence of any expression of the ORC1 protein.

### Origin properties in the knockout cells

Origin selection in mammalian cells varies from cell-cycle to cell-cycle in the same cells and certainly between cell lines. Since we intended to compare origins in four cell lines, with different passage histories, we were not sure how much overlap we would see between origins in the four cells. The results were clear. With nearly 50% of the origins mapped by our experiments in HCT116 cells overlapping with the origins reported in the literature as Core origins, we know that our experiments are identifying *bona fide* origins. The surprising discovery is that the vast majority of the origins in the WT cells were seen in one or more of the three ORC subunit deletion cells. This suggests that (a) origin selection still occurs in the absence of all six subunits of ORC and (b) the ORC deletion cells do not use an entirely new set of origins to replicate and survive in the absence of ORC.

The origins seen in the ORC knockout cells were generally similar to the origins in WT cells regarding GC content and preference for promoters, CpG islands and G quadruplexes. Thus, the general features that promote origin selection, GC content, promoter, CpG islands and G quadruplexes continue to select origins in the absence of ORC. The higher GC skew in the origins of the ORC knockout cells will be discussed below.

### The origins that are activated preferentially in the absence of ORC

Because there were a few hundred origins that were upregulated in all three *ORCΔ* cells (compared to WT cells), we examined them closer to see whether they had any sequence properties that set them apart from the vast majority of origins. These origins had a higher GC skew and TA skew and a lower overlap with promoters, CpG islands, R loops and G quadruplexes than the origins that overlapped with those in WT cells. As mentioned above, promoters, CpG islands and G quadruplexes have been reported in multiple laboratories to favor origin selection(50–52,54,55,58,69–77), so the decrease in their enrichment in the origins upregulated in ORC-KO cells suggest that the knockout cells may use slightly different properties of DNA or chromatin to select origins. Indeed, the increase in GC skew, TA skew and unique enrichment of (ATCC)n simple repeats in CommonKO_up set could suggest that these features help to license or fire origins in the absence of ORC, but of course, this is a correlation analysis and the association may not be causative.

In bacteria and in eukaryotes, it has been reported that the count of G exceeds that of C (GC skew positive) and T over A (TA skew positive) on the leading strand (78–81) and this trend peaks near origins (where the leading strand-lagging strand switch occurs). This increased GC and TA skew near origins has been suggested to be an evolutionary footprint arising from deamination of cytosines on the lagging strand template that is more often left single-stranded during DNA replication. The higher GC skew in the origins that are up-regulated in the ORC knockout cells either suggests that these origins have a longer evolutionary history or that increased GC skew favors MCM2-7 loading in the absence of ORC.

(ATCC)n repeats can on the opposite strand be represented as (TGGA)n repeats. It is thus very interesting that (TGGA)n repeats have been reported to be reluctant to assemble nucleosomes in *in vitro* selection experiments with a nearly 50% decrease in affinity for histone octamers compared to average DNA (82). Thus, one reason why these sequences promote origin function in the ORC knockout cells could be because they are less chromatinized and so more susceptible to MCM2-7 loading.

### None of the ORC subcomplexes on chromatin are better than the others at origin specification

There was a gradient of loading of ORC subunits in the knockout cells. The *ORC2Δ* cells had a great decrease of 4 subunits (ORC2, ORC3, ORC4 and ORC5) and a moderate decrease of 1 subunit (ORC1) on chromatin. The *ORC5Δ* cells had a great decrease of 2 subunits (ORC4 and ORC5) on chromatin. The *ORC1Δ* cells showed a decrease of only 1 subunit (ORC1) on chromatin. Our results show that if the remaining subunits formed subcomplexes that can assist in MCM2-7 loading, they are not much worse than the holocomplex for origin selection. One formal possibility is that ORC1 on its own and ORC2-5 on the other side can independently load MCM2-7 in unusual circumstances when the entire six-subunit ORC is absent. However, our current understanding of ORC structure and function makes this unlikely (2,3,18,19), and we have experimentally determined that hepatocytes without both ORC1 and ORC2 are still capable of loading functional MCM2-7 leading to normal liver development, liver regeneration, endo-reduplication and DNA synthesis (83).

### Functional and excess MCM2-7 loading

The ability of the ORC knockout cells to load functional MCM2-7 in excess (for dormant origin function), repeatedly (for re-replication), and at nearly the same kinetics in G1 as in WT cells highlights that these cells have a robust mechanism to load the MCM2-7 double hexamers in the virtual absence of ORC. Till now it could be argued that equal levels of MCM2-7 *association* with chromatin in the knockout cells compared to WT cells does not necessarily equate with comparable levels of *functional loading* of MCM2-7 double hexamers around the DNA. It could also be argued that an MCM2-7 loading system inactivated by the loss of ORC could barely load enough of the hexamers to permit the one round of replication required for cell cycle and cell proliferation. However, the results with hydroxyurea stressed cells or with stabilized Cdt1, suggest that the MCM2-7 loading machinery is still capable of loading excess functional MCM2-7 and/or repeatedly loading functional MCM2-7 in the same cell-cycle despite the absence of ORC.

### ORC-independent MCM2-7 loading

The near normal behavior of the cancer cells in the absence of the six-subunit ORC is a puzzle. In *S. cerevisiae*, ORC binding to the DNA causes it to bend, but it is not clear whether this bending is essential for the subsequent MCM2- 7 loading (84). Structural studies on yeast MCM2-7 hexamer complexed with CDT1 and on human MCM2-7 hexamer show that the MCM ring has an open gap between MCM2 and MCM5 (85–87). The size of the gap for human MCM2-7 has been estimated to be 10-15 Angstroms, which is too narrow to admit the 20 Angstrom diameter DNA double helix, but ORC is not needed to separate the MCM2 and 5 subunits to create the gap. Thus, there may be other ways, besides ORC, to either widen the gap to load a double-stranded DNA or to take advantage of naturally occurring denatured parts of DNA to let the MCM2-7 ring load on to a single strand of DNA. In addition, while the current understanding of yeast MCM2-7 loading is that ORC is required to load two MCM2-7 hexamers sequentially on DNA to produce the functional head-to-head double hexamer, it has been shown recently that single human MCM2- 7 hexamers loaded on DNA independently can come together to form a double hexamer of MCM2-7 (67). We have now ruled out that the origins used in these cells are very non-specific or very different compared to WT cells. We have also ruled out that the MCM2-7 loading in these cells is not robust and barely sufficient to carry out one round of DNA replication. The results suggest that while ORC may be essential to load MCM2-7 in many physiological conditions, during development, and even in current *in vitro* MCM2-7 loading reactions, the mammalian cancer cell is capable of loading functional and excess MCM2-7 in the absence of the six-subunit ORC. In this context, it is also very intriguing that certain ancient species of budding yeast in the *Hanseniaspora* lineage retain the *ORC1*, *ORC2*, *ORC4* and *ORC5* genes, but lack the *ORC3* and *ORC6* genes (88).

### Epigenetic regulation of origin firing

A strong role of epigenetics in origin specification in eukaryotic cells has been reported multiple times (89–93), particularly in view of the enrichment of origins on euchromatin and near promoters, but it has always been assumed that the chromatin decompaction at those sites help recruit ORC to those sites, which then leads to MCM2-7 loading. The fact that those epigenetically open sites are still the preferred sites of origins even in the absence of ORC and that the origins fired in the *ORCΔ* cells are still enriched in the early replicating (euchromatic) parts of the genome makes it likely that the open chromatin is also important for directly enabling MCM2-7 loading and even perhaps origin firing in the absence of ORC.

## Data availability

The raw data for SNS-seq have been deposited in NCBI’s Gene Expression Omnibus (GEO) and are accessible through GEO Series accession number GSE278065.

## ACKNOWLEDGEMENT

We thank all members of the Dutta and Pavri labs for helpful discussions and for Dr. Daniel Lew (MIT) for bringing the *Hanseniaspora* species to our attention.

## Author Contributions

MP did the SNS-seq experiments and DM did the initial mapping, YS did all the data analysis, ES and YS did the experiments in Figures 1, 8 and 9. YS and AD designed the project and wrote the paper. AD and RP supervised the experiments.

