## supplementary figures for "Specific origin selection and excess functional MCM2-7 loading in ORC-deficient cells"

**Supplementary figure S1** Permutation test results of colocalization between various origin peaks and simple repeat regions with TACC motif.

**Supplementary figure S2** ORC2 WB in ORC2 KO cells. A**)** Immunoblot of ORC2 in ORC3 IP in ORC2KO cell line.  B**)** Immunoblot of ORC2 in ORC2 KO cells with different ORC2 antibodies

**Supplementary Table S1** Adjusted p-values and 95% confident intervals of Welch's t-tests (**Fig. 5A, 6 and 7**).

**Supplementary Table S2** Adjusted p-values and effect sizes of comparison tests (**Fig. 4**). Bold:  Effect size > 0.15 or <-0.15

**Supplementary Table S3** Adjusted p-values of the Kolmogorov-Smirnov test, and Wasserstein distance (**Fig. 5B-D**).

**Supplementary Table S4** MCM2 Fluorescence signals in G1-MCM$^{DNA}$ positive cells used for Ergodic Rate Analysis calculation.  Each value is fluorescence intensity for MCM2 in one cell. There are 3 biological replicates for each cell line.

**Supplementary Fig. S1**

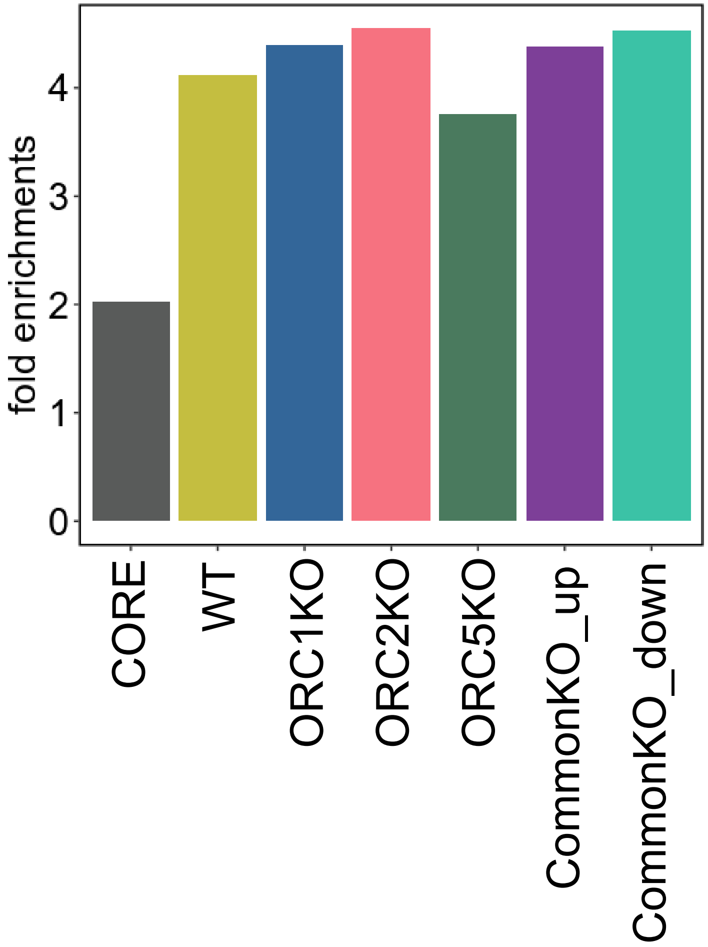

### Supplementary Fig. S2

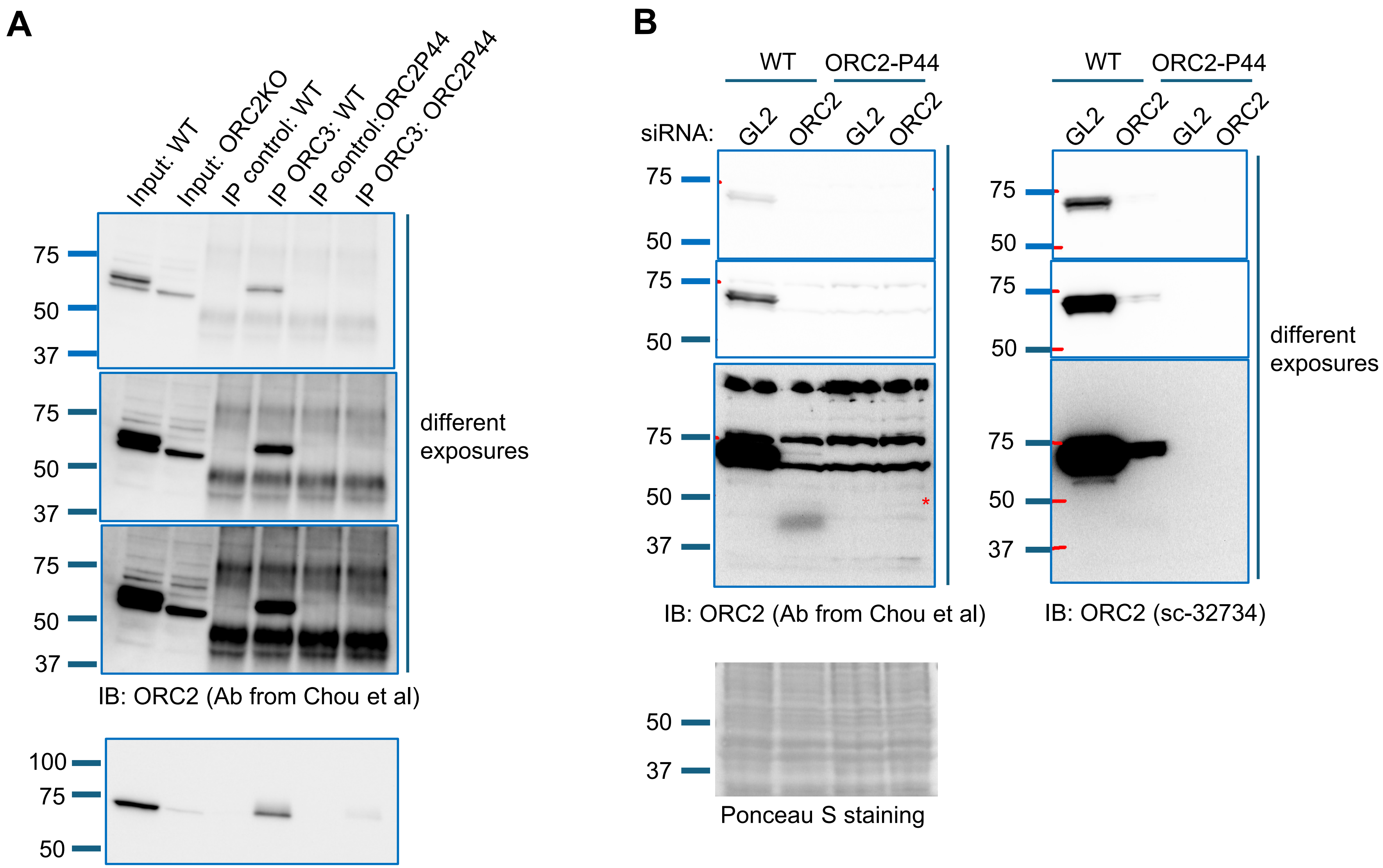

IB: ORC2 (Ab from Chou et al)

IB: ORC3

IB: ORC2 (Ab from Chou et al)

IB: ORC2 (sc-32734)

Ponceau S staining
